# Impact of Molecular Dynamics of Polyrotaxanes on Chondrocytes in Double Network Supramolecular Hydrogels under Physiological Thermomechanical Stimulation

**DOI:** 10.1101/2023.09.01.555869

**Authors:** Theofanis Stampoultzis, Vijay Kumar Rana, Yanheng Guo, Dominique P. Pioletti

## Abstract

Hyaline cartilage, a soft tissue enriched with a dynamic extracellular matrix, manifests as a supramolecular system within load-bearing joints. At the same time, the challenge of cartilage repair through tissue engineering lies in replicating intricate cellular-matrix interactions. This study attempts to investigate chondrocyte responses within double-network supramolecular hydrogels, tailored to mimic the dynamic molecular nature of hyaline cartilage. To this end, we infused non-covalent host-guest polyrotaxanes, by blending α-cyclodextrins as host molecules and polyethylene glycol as guests, into a gelatin-based covalent matrix, thereby enhancing its dynamic characteristics. Subsequently, chondrocytes were seeded into these hydrogels to systematically probe the effects of varied concentrations of introduced polyrotaxanes (instilling different levels of supramolecular dynamism in the hydrogel systems) on cellular responsiveness. Our findings unveiled an augmented level of cellular mechanosensitivity for supramolecular hydrogels compared to pure covalent-based systems. This is demonstrated by an increased mRNA expression of ion channels (TREK1, TRPV4, PIEZO1), signaling molecules (SOX9) and matrix remodeling enzymes (LOXL2). Such outcomes were further elevated upon external application of biomimetic thermomechanical loading that brought a stark increase in the accumulation of sulfated glycosaminoglycans and collagen. Overall, we found that matrix adaptability plays a pivotal role in modulating chondrocyte responses within double-network supramolecular hydrogels. These findings hold potential for advancing cartilage engineering within load-bearing joints.

**Graphical Abstract:** Double network supramolecular hydrogels enhanced chondro-induction significantly compared to purely covalent-based hydrogels as indicated by enhanced accumulation of extracellular matrix molecules.

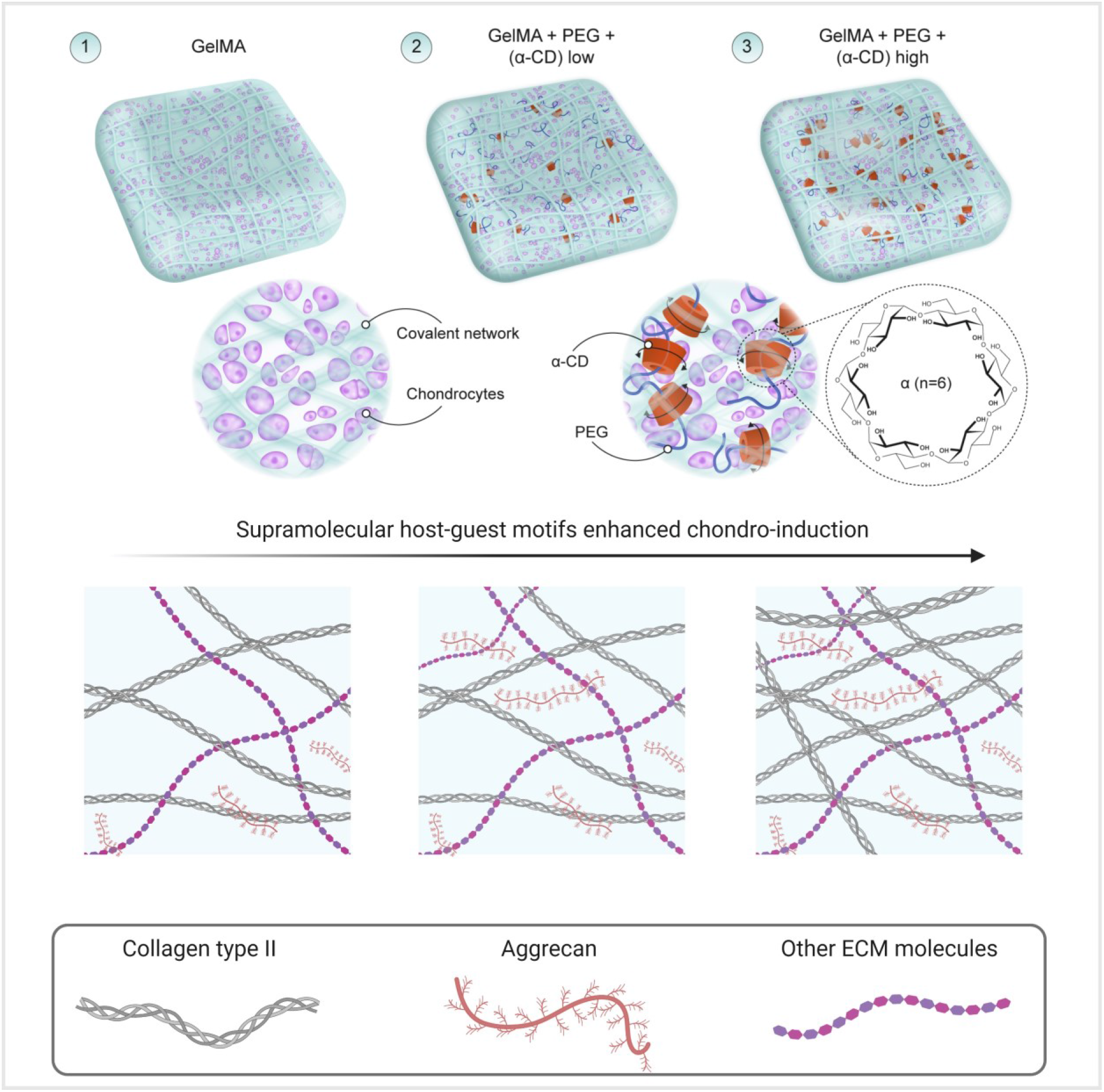

## 1. Introduction

In articular cartilage, chondrocytes and their dynamic local microenvironment constantly interact and communicate through biophysical and biochemical cues to regulate and guide various cell behaviors, such as cell differentiation.^1^ It is now globally accepted that dynamic temporal interactions predominantly mediated by chondrocyte adhesion to extracellular matrix and applied biomechanical stimuli, present a crucial role in transferring forces to and between cells that ultimately control chondrocyte function and tissue homeostasis.^2^ It is assumed that such interactions can be leveraged to alter cartilage disease and directly promote regeneration. Investigating the mechanisms by which physical cues and the nature of the cellular microenvironment are sensed by chondrocytes and how these are converted into biochemical signals is believed to be the gatekeeper to understand cartilage mechanobiology.^3^

Hydrogels are deemed highly attractive candidate materials to study how physical cues affect the chondrocyte responses *in vitro*, due to their inherent simplicity in terms of starting constituents and preparation, allowing for precise control of their chemical and physical properties.^4^ Nonetheless, thus far, previous studies have primarily focused on utilizing hydrogels crosslinked through covalent bonds to enhance compression resistance in case of applications for articulating joints. Despite their merits, covalent bonds tend to confine the synthesis of the extracellular matrix predominantly within the pericellular space.^5^ Conversely, biological tissues predominantly embrace the dominance of non-covalent interactions, presenting a stark divergence from the primarily covalent crosslinking strategies often observed in hydrogel investigations.^6^ In such regard, there is a growing interest in dynamic cell-adaptable hydrogels that can adjust and reorganize in response to mechanical stress or strain. As one example, supramolecular host-guest hydrogels are well-suited to study cartilage mechanobiology where the reversibility of the crosslinks occurs under physiological conditions.^7^ Supramolecular chemistry is entrenched in the rational design of specific and reversible molecular recognition motifs capitalizing on dynamic non-covalent interactions to create organized systems.^8^ Because of the dynamic nature of non-covalent interactions, supramolecular materials can rapidly respond to multifarious external stimuli, thereby recreating aspects of the dynamics present in living systems; making them a suitable candidate for cartilage studies.^9^

Although covalent hydrogels can be engineered to possess elastic and/or viscoelastic mechanical properties,^10^ they fail to replicate the inherent dynamics of the extracellular matrix found in tissues like hyaline cartilage. Herein, we have devised an approach that combines a covalent-based hydrogel with supramolecular polyrotaxane motifs, aiming to capitalize on the beneficial properties of supramolecular host-guest interactions (dynamic reversibility) within the covalent network. This integration enables the creation of a more permissive environment for encapsulated cells to interact with the hydrogel matrix during externally applied deformation, while simultaneously upholding the desirable characteristics, such as robustness, inherent in covalent bonding. By incorporating supramolecular polyrotaxanes, we enable the hydrogel to undergo reorganization and adaptability, mimicking the natural behavior of the extracellular matrix in hyaline cartilage.

So far despite its significance, a link between the temporal hierarchy of gel dynamics and adult chondrocyte behavior, particularly in response to externally applied biomimetic stimulation, remains elusive. Understanding how mechanobiological signals affect chondrocyte behavior is crucial for enhancing the outcomes of tissue engineering approaches.^11^ Thus, we further sought to examine mechanobiological interactions among dynamic matrix characteristics upon externally applied biomimetic thermomechanical load using host-guest supramolecular hydrogels. To the best of our knowledge this research work represents the first and only study to explore interactive effects between dynamic reversible crosslinks (polyrotaxanes) and physiologically relevant biomimetic thermomechanical loading for cartilage tissue engineering.

To this end, primary human chondrocytes encapsulated in supramolecular hydrogels were maintained in free swelling condition (static) as well as subjected to a long-term (up to 21 days) culture *via* a custom-made bioreactor apparatus designed to simulate transient thermomechanical stimuli as experienced in knee joint.^12^ PCR analysis was employed to investigate the early transcriptional interactions among hydrogels with static and dynamic host-guest crosslinks (polyrotaxanes), revealing significant transcriptional changes between the experimental groups in free swelling condition. Rates of biosynthesis were also analyzed by quantifying the deposition of sulfated glycosaminoglycans (sGAGs) and total collagen type following thermomechanical stimulation. Histological analysis was further utilized to visualize and detect the spatial distribution of these molecules. Overall, this study underscores the significance of a dynamic extracellular matrix (ECM) akin to that found in native cartilage, accentuating how mechanobiological cues intricately guide chondrocyte biosynthetic responses within the dynamic hydrogel milieu.

## 2. Materials & Methods

α-Cyclodextrin was purchased from Sigma Aldrich (C4642, ≥98%). Polyethylene glycol 2000 was purchased from Sigma Aldrich (Mn = 2000 Da, for synthesis). Gelatin Type A from porcine skin (ref. G2500) and methacrylic anhydride (ref. 276685) were purchased from Sigma Aldrich.

### 2.1 Preparation of host-guest molecules and supramolecular hydrogels

Host-guest supramolecular complexes were prepared by dissolving two different concentrations of alpha cyclodextrin (α-CD) (12 mg and 36 mg) in a PEG/PBS solution (6.5% wt). The mixture was thoroughly mixed and allowed to equilibrate for a specific period (1 day prior to experiments) to ensure the formation of the complexes. Methacrylated gelatin (GelMA) was synthesized as previously described^13^ and then dissolved in the [PEG-α-CD]/PBS system at a final concentration of 7% wt. The solution was gently stirred and heated at 37 °C until complete dissolution of GelMA was achieved. For the preparation of non-supramolecular hydrogels, GelMA was dissolved in PBS at a final concentration of 7% wt.

### 2.2 NMR analysis and Isothermal Titration Calorimetry (ITC)

^1^H-NMR spectra were acquired using a Bruker Avance NMR spectrometer (400 MHz) with a BBI probe and processed with MestReNOVA software, as previously described.^11^ Chemical shifts were reported in ppm, rounded to the nearest 0.01 ppm for 1H NMR.

ITC experiments were conducted in phosphate buffered saline (PBS) at 298.15 K using the MicroCal PEAQ-ITC. The sample cell contained the guest PEG2k solution, and the injection syringe contained the host alpha-CD solution. The concentrations of both alpha-CD and PEG2K were calibrated using ITC titration in a blank PBS solution. Heat evolution was observed when a specific concentration of host or guest was titrated into the blank PBS solution or vice versa to know the correct concentration for the final measurements.

### 2.3 Experimental groups

This study encompassed three distinct phases. In Phase I, the objective was to assess the effects of incorporating dynamic host-guest polyrotaxanes into a covalent network and their subsequent impact on chondroinduction, particularly on the expression of key chondrogenic genes. This evaluation involved a comparative analysis with motif-absent (single network) hydrogels under free-swelling conditions. Advancing to Phase II, the focus shifted to the identification of potential enhancements observed in Phase I, now at the protein level. Subsequently, Phase III involved subjecting the hydrogels to biomimetic thermomechanical stimulation under hypoxia. The overarching goal was to enhance chondrocyte biosynthesis by more accurately mimicking cartilage *in vivo* milieu. Phase I and Phase II were executed over a relatively condensed timeframe (Day 16), while Phase III extended over a more extended duration (Day 21).

### 2.4 Cell expansion, encapsulation and bioreactor culture

Primary human chondrocytes were derived from a 22-year old male donor (P10970 Innoprot, Spain). Cells were expanded in T-75 culture flasks inside chondrocyte basal medium (Alpha minimum essential medium (α-MEM)), supplemented with 10% fetal bovine serum (FBS), 1% L-glutamine, 10 mM 4-(2-hydroxyethyl)-1-piperazineethanesulfonic acid (HEPES), 10 mM nonessential amino acids (NEEA), 1% penicillin, 1% streptomycin, 5 ng/mL fibroblast growth factor (FGF) and 1 ng/mL human recombinant transforming growth factor beta 1 (TGF-β1), up to passage 4.

Hydrogel precursor solution were prepared, as described earlier, inside phosphate buffered saline (PBS) containing LAP photoinitiator (Lithium phenyl-2,4,6-trimethylbenzoylphosphinate, at a final concentration of 0.1 mg/ml). Passage 4 chondrocytes, at a seeding density of 10^7^ cells per ml, were resuspended in the hydrogel precursor and carefully pipetted into a custom-designed mold. The chondrocyte/hydrogel suspension was then crosslinked using a 405 nm wavelength light source for 2 minutes. The resulting constructs were cultured in a differentiation medium composed of FBS-free Dulbecco’s Modified Eagle’s medium supplemented with insulin–transferrin–selenium (ITS-IV, 10%), L-ascorbic acid (VC, 1%), 10 ng/mL TGF-β1, and other additives (10 mM HEPES and 10 mM NEAA). The cell-seeded constructs were prepared as a batch and subsequently randomly distributed among different study groups. Following the seeding step, all samples were pre-cultured for 7 days in a cell growth medium within standard incubators (32°C, 5% CO2, 21% O2). Next, the constructs were transferred to a bioreactor culture system where intermittent biomimetic thermomechanical signals (32–39 °C, 5% pre-strain, 10% amplitude at 1 Hz every other day) and a low oxygen tension environment (4% O2) were applied until day 21. We replicated all forms of stimulation by utilizing a custom-made bioreactor that we developed in our laboratory.^12^ The compressive regime was specifically designed to imitate normal physical activity. To model the temperature increase resulting from cyclic compression, we applied curve-fitting based on *in vivo* data during jogging over the same 1.5-hour period and as previously described.^14^ After stimulation was ceased, constructs were allowed to recover within standard incubators (32°C, 5% CO_2_). Constructs were collected for analysis on day 16 and day 21.

### 2.5 Quantitative real time PCR

After a 16 and 21 days of bioreactor culture, both stimulated and non-stimulated samples were promptly immersed in 0.3 ml ice-cold TRIzol (Invitrogen) and stored at -80 °C for subsequent RNA isolation. To prepare the samples, TRIzol was added to each sample on ice, followed by vigorous homogenization. Subsequently, 0.1 ml of chloroform was added, and the mixture was hand-shaken for 15 seconds and then centrifuged at 4 °C for 10 minutes. The aqueous layer, containing the RNA, was carefully collected and combined with an equal volume of 70% ethanol through pipetting. RNA isolation was performed using the Nucleospin XS kit according to the manufacturer’s instructions. The isolated RNA was quantified using a NanoDrop 1000, and then reverse-transcribed into cDNA using the Taqman Reverse Transcription Reagents (Applied Biosystems) in a 50 μL reaction volume as previously described.^15^ The reaction mixture included the master mix, random hexamer, and RNA sample.

For qRT-PCR analysis, the Fast SYBR Green PCR Master Mix (Applied Biosystems) was employed in a final volume of 20 μL. Each reaction contained 1 μL of synthesized cDNA. The target genes selected for qRT-PCR included background potassium channel (TREK1), lysyl oxidase-like 2 (LOXL2), aggrecan (ACAN), sex-determining region Y-type (SRY, SOX9), Piezo type mechanosensitive ion channel component 1 (PIEZO1), transient receptor potential cation channel subfamily V member 4 (TRPV4) and collagen type II (COL2A). To ensure accurate normalization, Ribosomal protein L13a (RPL13A) was utilized as the housekeeping gene. The relative gene expression of the experimental samples was determined using the ΔΔCT method. Primers were synthesized by Microsynth (Balgach, Switzerland).

### 2.6 Sample preparation for histology

To assess the production and distribution of the extracellular matrix (ECM), we employed histology. After 16 and 21 days, cell-hydrogel samples were harvested, and the chondrocytes were fixed in 4% paraformaldehyde overnight at 4 degrees Celsius. The following day, the samples were sequentially transferred to 15% and 30% sucrose solutions (Sigma) before being snap frozen in optimal cutting temperature compound (Sakura Tissue-Tek). Subsequently, the samples were sectioned into 7-μm thick slices using a standard cryostat. Imaging of histologic slides from different samples was performed using the tile scan method with a 20× magnification on an Olympus VS120 whole slide scanner. Throughout the imaging process, the laser intensity and exposure duration were consistently set for all samples to ensure accurate comparisons.

### 2.7 Mechanical characterization of hydrogels

Cylindrical specimens with dimensions Ø: 6 mm and h: 3 mm were submerged in bidistilled water and underwent unconfined compression experiments. The experiments were conducted using an Electropuls Dynamic Test System (Instron E3000, Instron, Norwood, Massachusetts, USA) at room temperature, with a 5% prestrain and a 1 Hz sinusoidal load of 10% amplitude. To determine the level of energy dissipation, the area enclosed by the hysteresis loop was measured and subsequently normalized to the volume of each sample.

### 2.8 Statistical analysis

Statistical analysis was conducted using analysis of variance (ANOVA) followed by Tukey’s post hoc tests for comparing mechanical data (n = 5) and gene expression data (n = 4) in multiple group comparisons. The results are presented as mean + standard deviation. The significance levels are denoted as (*) for p ≤ 0.05, (**) for p ≤ 0.01, and (***) for p ≤ 0.001. Statistical computations were performed using Origin Pro 2021 software. Additionally, to validate gene expression patterns, two independent experiments were conducted each of at least 3 biological replicates.

## 3. Results

### 3.1 Inclusion complex formation of linear polyethylene glycol (PEG) and *α*-cyclodextrin (*α*-CD)

Figure 1a shows a schematic illustration of typical polyrotaxanes consisting of a linear polymer chain (PEG) of 2kDa molecular weight and threaded rings (α-CD). The formation of inclusion complexes in an aqueous solution was confirmed and monitored by isothermal titration calorimetry (ITC). Figure 1b shows a typical ITC sigmoid thermogram indicating stable complex formation, while the stoichiometric ratio was close to 2.

**Figure 1.**
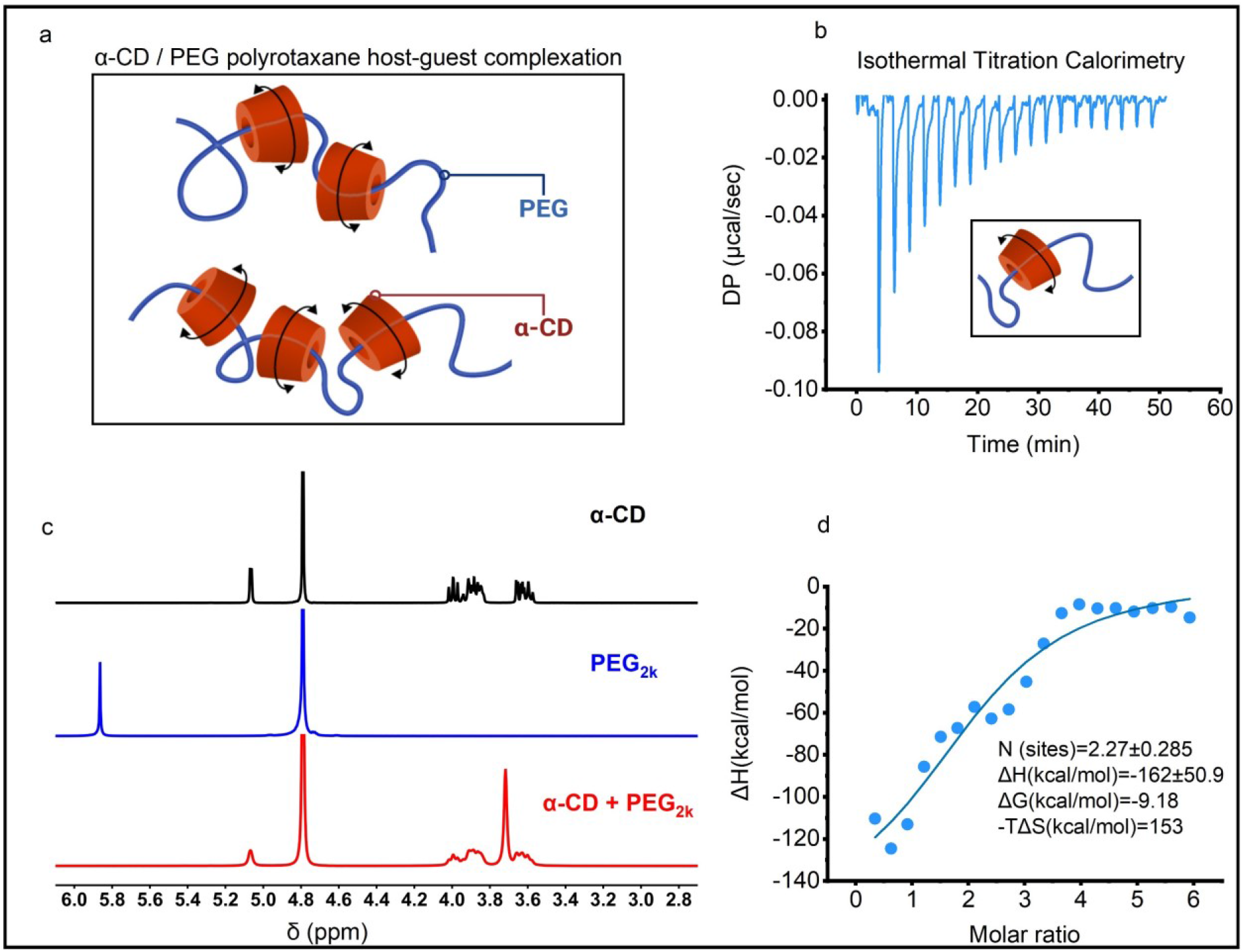
a) Schematic illustration of elemental motion of polyrotaxanes, b) measured heat of interaction obtained from ITC from sequential α-CD injections into an PEG2k solution, c) representative ^1^H NMR spectra of free α-CD (black line), free PEG2k (blue line), and α-CD + PEG2k complexes (red line), d) fitting of heat mathematical integration as a function of molar ratio for the PEG2k / α-CD inclusion complex.

^1^H NMR spectroscopy was further utilized to investigate the relative position of the host-guest complexation between α-CD (a host) and PEG2k chain (a guest). Figure 1c shows the NMR data of free α-CD (black line), free PEG2k (blue line), and α-CD + PEG2k complexes (red line). It is evident that upon incorporation into the host, the protons of the PEG2k chain experienced a complete upfield shift (shielded), while the protons of the α-CD cage exhibited peak broadening compared to the free α-CD. This observation confirms the occurrence of host-guest complexation between α-CD and the PEG2k polymer chain.

### 3.2 Phase I: Dynamic polyrotaxane host-guest complexation altered gene categories encoding multiple signaling pathways in free-swelling hydrogels

Quantitative real-time PCR (qPCR) analysis was conducted on free-swelling hydrogels (labeled as 1-3) to discern the impact of dynamic host-guest complexation (polyrotaxanes) at transcriptional level on day 16 of culture.

The findings unveiled profound changes in the expression of genes implicated in crucial cellular signaling pathways (SRY-related HMG-box gene 9, SOX9), genes associated with matrix remodeling enzymes (Lysyl oxidase homologue 2, LOXL2), extracellular matrix proteins (including Aggrecan, ACAN) and thermo-mechano-regulated ion channels (including potassium and calcium transducers, TREK1, TRPV4, PIEZO1) as illustrated in figures 2a-f.

**Figure 2.**
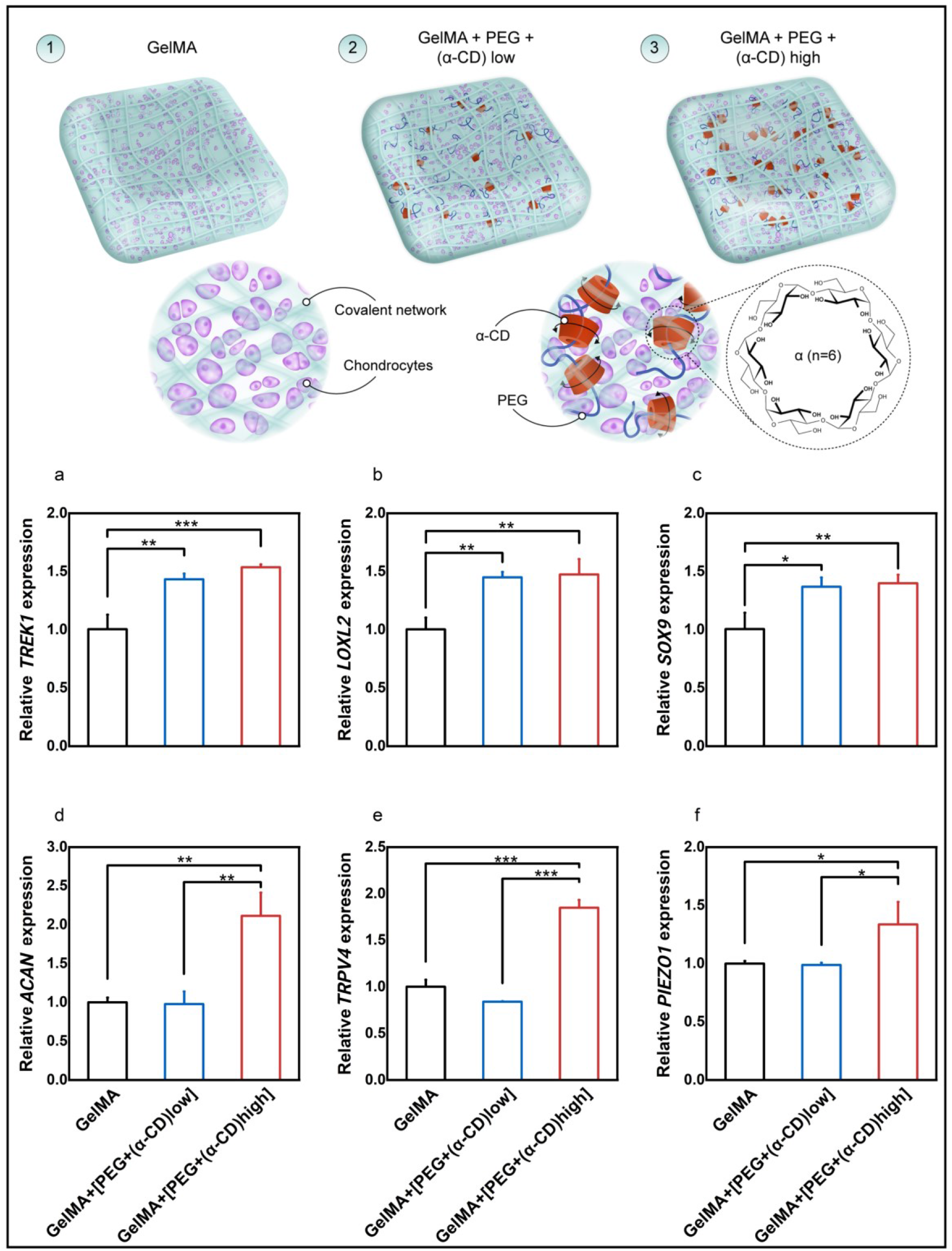
Schematic illustration depicting the various types of hydrogels employed in the study. The polyethylene glycol (PEG) concentration remained consistent across all supramolecular hydrogels, while the alpha cyclodextrin (α-CD) concentration increased progressively from left to right (a-f). Comparison of the relative expressions of genes of interest, with RPL13a serving as the housekeeping gene.

Overall, the inclusion of α-CD/PEG polyrotaxanes within the gelatin-based covalent network yielded mixed results worth noting. At lower concentrations, TREK1, LOXL2, and SOX9 genes displayed significant increases in expression, indicating a positive impact. However, the transcription plateaued with higher concentrations, not contributing to further improvements in these genes (Figure 2a-c). Conversely, other major chondrogenic genes such as ACAN, PIEZO1, and TRPV4 exhibited no discernible changes at lower concentrations. Yet, an astonishing revelation emerged; these three genes demonstrated a synchronized response pattern, undergoing a remarkable increase in expression at higher concentrations spanning from 30 to 200% (Figure 2c-f).

These compelling results highlight the transformative influence of dynamic host-guest complexation on the gene expression profile, underscoring the potential of this approach in modulating vital cellular pathways involved in chondrocyte physiology.

### 3.3 Phase II: The impact of dynamic host-guest polyrotaxanes on sulfated proteoglycans and total collagen synthesis and distribution during free-swelling culture of hydrogels

In cartilage tissue engineering, scaffolds must provide mechanical support while facilitating the deposition of extracellular matrix (ECM) by embedded chondrocytes to promote the development of neo-cartilaginous tissue.^16,17^ In this series of experiments, we aimed to investigate the influence of double network supramolecular hydrogels on the uniform secretion of the cartilaginous matrix during free-swelling conditions (without external stimulation up to day 16).

Collagens, as fibrillar proteins, play a crucial role in determining the shape and microarchitecture of articular cartilage.^18^ Sulfated glycosaminoglycans (sGAGs), major components of proteoglycans such as aggrecan, contribute to water retention and provide compressive strength.^19^ We utilized histological staining techniques to examine the deposition of these key cartilage matrix molecules and assess the development of neo-cartilaginous tissue over time.

In agreement with the gene expression data, our findings have unveiled remarkable disparities in both collagen and sGAG content, intricately linked to the concentration of α-CD/PEG polyrotaxanes within the hydrogels. Astonishingly, higher concentrations of α-CD which translates to higher dynamism in the hydrogels system (due to higher level of α-CD/PEG complexations) have been discovered to foster a substantial accumulation of aggrecan, vividly exemplified by intensified alcian blue staining. Furthermore, the diverse compositions of the hydrogels have unraveled distinct patterns in total collagen deposition, with higher concentrations of α-CD showcasing an evident surge in overall collagen accumulation.

### 3.4 Phase III: Supramolecular host-guest complexation modulates the effects of biomimetic thermomechanical stimulation on chondrocyte biosynthesis in a dose-dependent manner

In this series of experiments, we extended the culture period from 16 days to 21 days while applying biomimetic thermomechanical stimulation upon hypoxia treatment to the chondrocyte-laden hydrogels. Notably, our findings reveal consistent and compelling trends in the total collagen and glycosaminoglycan (GAG) content. Increasing concentrations of α-CD within the covalent-based network (while keeping the PEG concentration the same) demonstrated a remarkable capacity to promote augmented collagen accumulation (as shown in Sirius Red and Masson’s Trichrome stainings), accompanied by significant deposition of sulfated GAGs (as shown in Alcian Blue and Safranin-O stainings) compared to pure covalent-based hydrogels. These outcomes underscore the pivotal role of host-guest supramolecular motifs in governing collagen and GAG secretion, thereby emphasizing the potential of double network supramolecular hydrogels, to facilitate the development of functional neo-cartilaginous tissue especially upon biomimetic thermomechanical stimulation.

**Figure 3.**
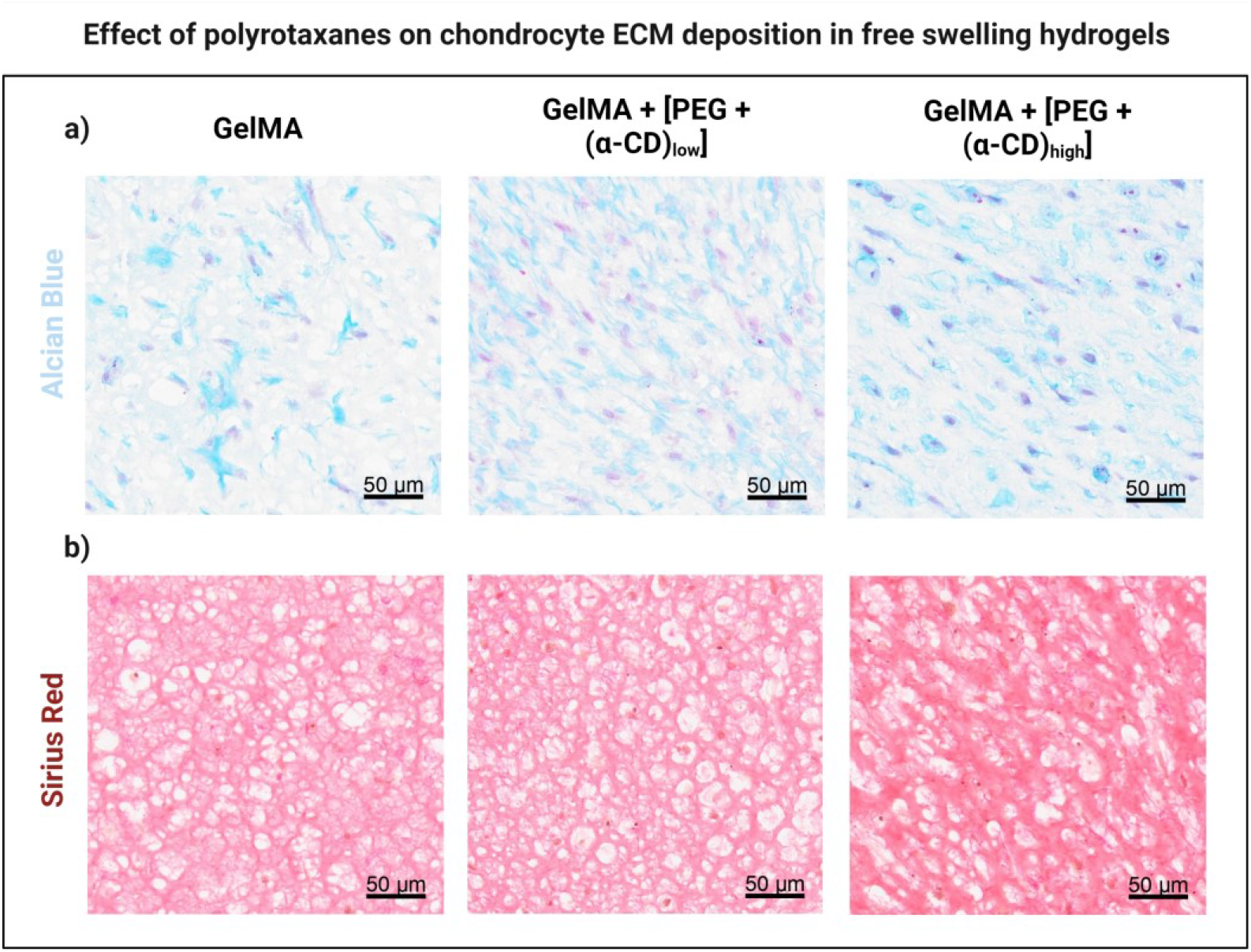
Histological analysis of neo-cartilage constructs under free-swelling conditions. a) representative images of Alcian Blue staining for sulfated glycosaminoglycan & glycoproteins content (blue), b) representative images of Sirius Red staining for general collagen content (red). The intensity of Alcian blue staining is noticeably higher in supramolecular hydrogels with a higher concentration of alpha cyclodextrin (α-CD) molecules. Additionally, Sirius Red staining reveals a localized increase as the concentration of α-CD molecules increases. Scale bar: 50 μm, objective 20x.

In addition to the histological stainings, we analyzed the mRNA transcription of aggrecan (ACAN, Figure 4e) and collagen type II (COL2A, Figure 4f) genes among the different hydrogels. The inter-group comparisons aligned with the histological findings, demonstrating a substantial upregulation of these pivotal genes, with an increase of 200-250%, respectively.

**Figure 4.**
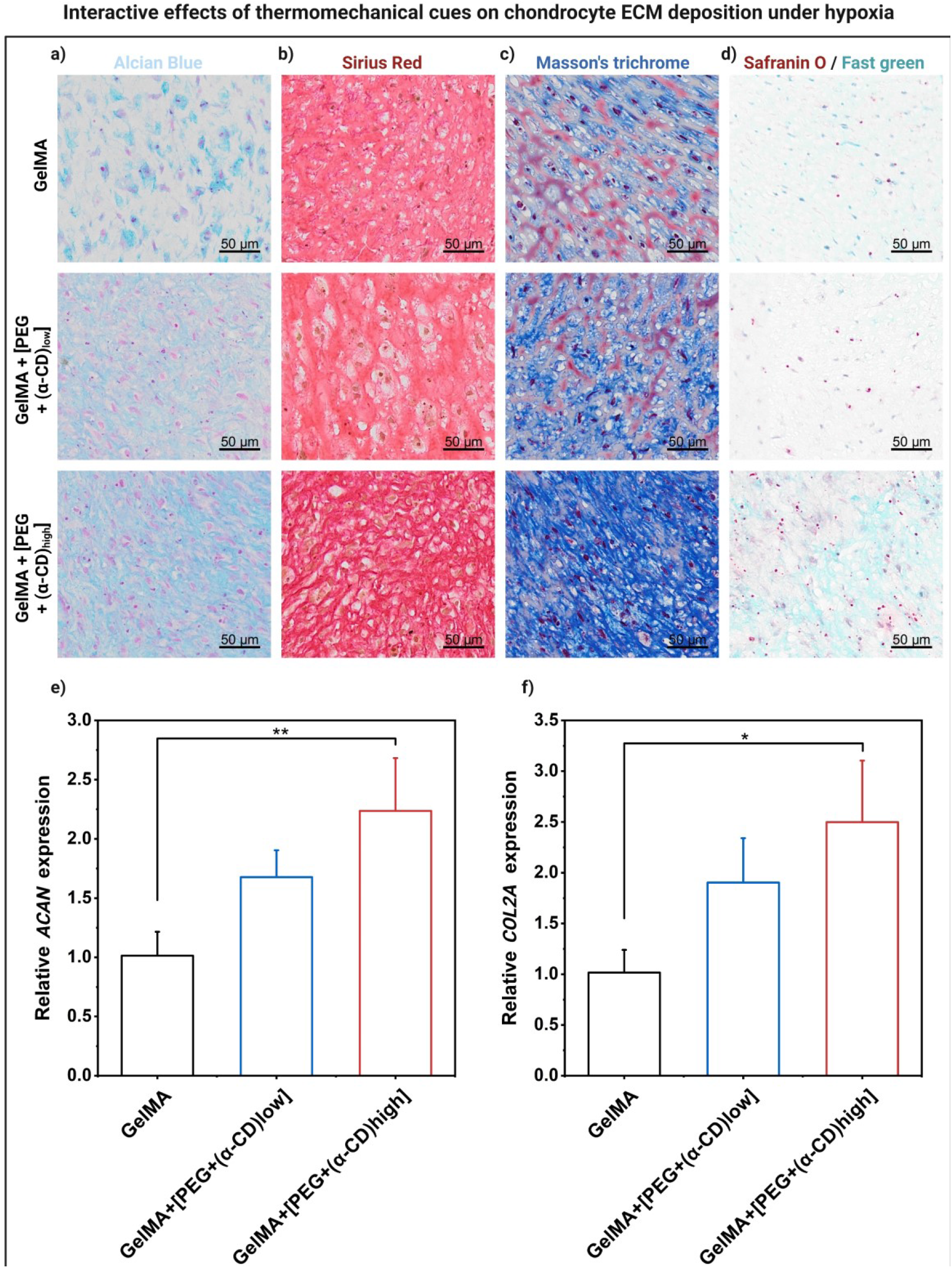
Interactive effects of mechanobiological cues on chondrocyte ECM deposition. a) representative images of Alcian Blue staining for sulfated glycosaminoglycan & glycoproteins content (blue), b) representative images of Sirius Red staining for general collagen content (red), c) representative images of Masson’s trichrome staining for total collagen content (blue) d) representative images of Safranin-O/Fast green staining for sulfated glycosaminoglycan content are shown for the different types of hydrogels after the last loading cycle was ceased on day 21. Application of biomimetic thermomechanical loading under hypoxia significantly enhanced cartilage-related matrix accumulation especially in the case of supramolecular hydrogels. <u>Alcian Blue</u>: sulfated GAGs and glycoproteins are stained blue, and the nuclei and cytoplasm pink, <u>Sirius Red</u>: collagen is stained red and the nuclei dark brown, <u>Masson’s trichrome</u>: collagen is stained blue, nuclei are stained dark brown, <u>Safranin-O/Fast green</u>: cartilage matrix will be stained orange to red, the nuclei will be stained black, and the background light green. Scale bar: 50 μm, objective 20x. e) Comparison of the relative expressions of aggrecan (ACAN) and f) collagen type II (COL2A) after the last loading cycle was ceased, with RPL13a serving as the housekeeping gene.

## 4. Discussion

The interdependence of chondrocytes and their extracellular matrix (ECM) is a fundamental aspect of cell behavior and function.^20^ To comprehend these relationships *in vitro*, hydrogels are a promising candidate due to the possibility of precisely controlling their chemical and physical properties. However, existing studies have predominantly utilized purely covalently crosslinked hydrogels, which fail to confer the essential dynamicity found in the ECM of native tissues.^21,22^ Supramolecular hydrogels offer a dynamic non-covalent alternative that more accurately reflects the native environment of hyaline cartilage.^23^ Utilizing supramolecular non-covalent motifs such as polyrotaxanes, this study aimed to engineer dynamic hydrogels with the ability to reorganize under free-swelling conditions. More specifically, we have developed a method that integrates a covalent-based hydrogel with supramolecular polyrotaxane patterns, with the intention of harnessing the advantageous attributes of supramolecular host-guest interactions (characterized by dynamic reversibility) within the covalent network. The addition of dynamic polyrotaxanes into the covalent network hydrogels was intended to enhance the biosynthetic capacity of neocartilage constructs. The underlying hypothesis suggested that by replicating the natural characteristics observed in cartilage extracellular matrix (e.g inherent dynamic network of molecules), chondrocytes would be prompted to produce essential matrix proteins. Subsequently, the investigation delves into the biophysical impacts of applied thermomechanical loads on chondrocytes encapsulated in these double network hydrogels. Given that native cartilage experiences transient thermal cues during joint loading upon a hypoxic environment, replicating these interactions *in vitro* became a focal point for assessing the potential acceleration of chondrocyte biosynthesis.

Through our rigorous investigation, we have uncovered a fascinating interplay between double network supramolecular hydrogels, thermomechanical loading, and hypoxic conditions in regulating chondrogenesis. Our research unveiled that when polyrotaxanes are incorporated into the covalent network, especially at increased concentrations, alongside thermomechanical loading and reduced oxygen tension, a significant increase in chondrogenic markers and cartilage-related proteins is observed. This synergistic effect suggests a unique interplay between these cues that holds significant promise in the field of cartilage tissue engineering, highlighting the importance of a multifaceted approach to this complex challenge.

Remarkably, our findings indicate a substantial augmentation in chondrogenesis within the double network supramolecular hydrogels, surpassing that observed in the covalently crosslinked (single network) hydrogel counterparts. Moreover, a discernible relationship emerges, whereby the extent of host-guest complexation (polyrotaxanes) exhibits a direct correlation with the magnitude of chondrogenic differentiation. This positive correlation is substantiated through quantifiable variations in mRNA expression levels of critical chondrogenic marker genes, namely SOX9, ACAN, and LOXL2. Such improvements were detected at protein level as well. These findings are consistent with previous research^24, 7, 25^ supporting the notion that supramolecular hydrogels, particularly those featuring stronger host-guest interactions, offer a promising platform for the development of tissue-engineered constructs with superior chondrogenic potential.

In our investigation, the hydrogel formulations were also subjected to bioreactor culture conditions that incorporated thermomechanical signals under low oxygen tension. Our previous studies have extensively elucidated the significance and relevance in replicating cartilage self-heating *in vitro* and demonstrated thoroughly how loading-induced evolved temperature can be harnessed *in vitro* to accelerate tissue maturation through expression of major structural proteins.^26^ Building upon our prior findings where load, heat and hypoxia interactions were studied, herein we aimed to introduce the inherent dynamic nature of the culture environment as an additional parameter in this multifaceted equation. The relevance of including PEG/alpha cyclodextrin-based polyrotaxanes into the robust covalent network stems from their capacity to partially replicate the dynamic interactions found in natural cartilage.^27,28^ Through the incorporation of dynamic polyrataxanes, we aim to mimic the innate non-covalent interactions inherent in cartilage tissue, playing a pivotal role in its mechanical composure and potential signals they transfer to chondrocytes. Our analysis of the resulting protein synthesis levels in the stimulated samples revealed that the double network hydrogels exhibited a notable upregulation of cartilage matrix proteins compared to the pure covalently crosslinked hydrogel.

Our findings strongly support the notion of the pivotal role of ion channels in mediating the effects of thermomechanical loading on chondrocytes. We identified three ion channels, namely TREK1, TRPV4, and PIEZO1, whose expression was significantly upregulated in the presence of host-guest reversible polyrotaxanes. While it could be argued that our channel analysis has only been performed in free-swelling constructs and not in stimulated hydrogels, we have previously shown that thermomechanical loading increases the expression of these channels at protein levels as well.^12,26^ Hence, we contend the validity of our hypothesis, suggesting that the observed effects likely arise from the interplay between various stimuli and the multifaceted responses of chondrocytes.

We also conducted a systematic investigation into the distinct influences stemming from the separate incorporation of PEG and alpha cyclodextrin moieties into the covalent network. Interestingly, we found no significant changes in the expression of major genes among the different experimental groups, indicating that the observed effect primarily arises from the complexation of alpha cyclodextrin and PEG (See supplementary figure 1). To ensure the validity of our observations, we diligently measured the bulk mechanical properties of the hydrogels, specifically focusing on energy dissipation levels before cell encapsulation (See supplementary figure 2). Encouragingly, our results demonstrated consistent mechanical attributes across the experimental groups (prior to cell encapsulation), suggesting that no other parameter significantly influenced the observed effects. These findings lend strong support to the relevance and reliability of our conclusions.

In conclusion, this study provides crucial insights into the dynamic nature of cartilage, even under free-swelling conditions, and the role of dynamic polyrotaxanes, thermomechanical loading, and hypoxia in regulating chondrogenesis. By incorporating supramolecular motifs into a covalent-based hydrogel system, we aimed to partially mimic the intricate dynamic and non-covalent interactions present in natural cartilage. Supramolecular hydrogels with high amount of host-guest crosslinks, when subjected to thermomechanical stimulation and hypoxic conditions, enhance chondrogenic markers and proteins. This highlights their potential as a promising platform for advanced tissue-engineered constructs with improved chondrogenic capabilities

## Acknowledgement

We thank Mrs. Sandra Jaccoud for her overall technical support in cryosection of the cell-laden hydrogels. We would like to thank the Histology Core Facility at EPFL for conducting the histological stainings. Bottom part of Graphical abstract was ‘‘created with BioRender.com’’ for which we have the license for publication. This research project is supported by a SNF grant # CRSII5_189913.

## References

1. Michelacci, Y.M., Baccarin, R.Y.A., and Rodrigues, N.N.P. (2023). Chondrocyte Homeostasis and Differentiation: Transcriptional Control and Signaling in Healthy and Osteoarthritic Conditions. Life 13, 1460. 10.3390/life13071460.

2. Gao, Y., Liu, S., Huang, J., Guo, W., Chen, J., Zhang, L., Zhao, B., Peng, J., Wang, A., Wang, Y., et al. (2014). The ECM-cell interaction of cartilage extracellular matrix on chondrocytes. Biomed Res. Int. 2014. 10.1155/2014/648459.

3. Conor, C.J.O., Case, N., and Guilak, F. (2013). Mechanical regulation of chondrogenesis. Stem Cell Res. Ther. 4, 1. 10.1186/scrt211.

4. Stampoultzis, T., Karami, P., and Pioletti, D.P. (2021). Thoughts on cartilage tissue engineering: A 21st century perspective. Curr. Res. Transl. Med. 69, 103299. 10.1016/j.retram.2021.103299.

5. Richardson, B.M., Walker, C.J., Maples, M.M., Randolph, M.A., Bryant, S.J., and Anseth, K.S. (2021). Mechanobiological Interactions between Dynamic Compressive Loading and Viscoelasticity on Chondrocytes in Hydrazone Covalent Adaptable Networks for Cartilage Tissue Engineering. Adv. Healthc. Mater. 10. 10.1002/adhm.202002030.

6. Gottardi, R., Hansen, U., Raiteri, R., Loparic, M., Düggelin, M., Mathys, D., Friederich, N.F., Bruckner, P., and Stolz, M. (2016). Supramolecular organization of collagen fibrils in healthy and osteoarthritic human knee and hip joint cartilage. PLoS One 11, 1–13. 10.1371/journal.pone.0163552.

7. Yang, R., Xue, W., Ma, X., Ren, Y., Xu, L., Kong, W., Zhang, W., Wang, P., Tan, X., and Chi, B. (2023). Engineering the dynamics of biophysical cues in supramolecular hydrogels to facile control stem cell chondrogenesis for cartilage regeneration. Compos. Part B Eng. 250, 110429. 10.1016/j.compositesb.2022.110429.

8. Seiffert, S., and Sprakel, J. (2012). Physical chemistry of supramolecular polymer networks. Chem. Soc. Rev. 41, 909–930. 10.1039/c1cs15191f.

9. Webber, M.J., Appel, E.A., Meijer, E.W., and Langer, R. (2015). Supramolecular biomaterials. Nat. Mater. 15, 13–26. 10.1038/nmat4474.

10. Roberts, J.J., Earnshaw, A., Ferguson, V.L., and Bryant, S.J. (2011). Comparative study of the viscoelastic mechanical behavior of agarose and poly(ethylene glycol) hydrogels. J. Biomed. Mater. Res. - Part B Appl. Biomater. 99 B, 158–169. 10.1002/jbm.b.31883.

11. O’Conor, C.J., Leddy, H.A., Benefield, H.C., Liedtke, W.B., and Guilak, F. (2014). TRPV4-mediated mechanotransduction regulates the metabolic response of chondrocytes to dynamic loading. Proc. Natl. Acad. Sci. U. S. A. 111, 1316–1321. 10.1073/pnas.1319569111.

12. Nasrollahzadeh, N., Karami, P., Wang, J., Bagheri, L., Guo, Y., Abdel-Sayed, P., Laurent-Applegate, L., and Pioletti, D.P. (2022). Temperature evolution following joint loading promotes chondrogenesis by synergistic cues via calcium signaling. Elife 11, 1–23. 10.7554/eLife.72068.

13. Karami, P., Nasrollahzadeh, N., Wyss, C., O’Sullivan, A., Broome, M., Procter, P., Bourban, P.E., Moser, C., and Pioletti, D.P. (2021). An Intrinsically-Adhesive Family of Injectable and Photo-Curable Hydrogels with Functional Physicochemical Performance for Regenerative Medicine. Macromol. Rapid Commun. 42. 10.1002/marc.202000660.

14. Becher, C., Springer, J., Feil, S., Cerulli, G., and Paessler, H.H. (2008). Intra-articular temperatures of the knee in sports - An in-vivo study of jogging and alpine skiing. BMC Musculoskelet. Disord. 9, 1–7. 10.1186/1471-2474-9-46.

15. Nasrollahzadeh, N., Karami, P., and Pioletti, D.P. (2019). Control of Dissipation Sources: A Central Aspect for Enhancing the Mechanical and Mechanobiological Performances of Hydrogels. ACS Appl. Mater. Interfaces 11, 39662–39671. 10.1021/acsami.9b15450.

16. Akalp, U., Bryant, S.J., and Vernerey, F.J. (2016). Tuning tissue growth with scaffold degradation in enzyme-sensitive hydrogels: A mathematical model. Soft Matter 12, 7505–7520. 10.1039/c6sm00583g.

17. Huey, D.J., Hu, J.C., and Athanasiou, K.A. (2012). Unlike bone, cartilage regeneration remains elusive. Science (80-.). 338, 917–921. 10.1126/science.1222454.

18. Sophia Fox, A.J., Bedi, A., and Rodeo, S.A. (2009). The basic science of articular cartilage: Structure, composition, and function. Sports Health 1, 461–468. 10.1177/1941738109350438.

19. Roughley, P.J. (2006). The structure and function of cartilage proteoglycans. Eur. Cells Mater. 12, 92–101. 10.22203/eCM.v012a11.

20. Anderson, D.E., and Johnstone, B. (2017). Dynamic mechanical compression of chondrocytes for tissue engineering: A critical review. Front. Bioeng. Biotechnol. 5, 1–20. 10.3389/fbioe.2017.00076.

21. Nicodemus, G.D., Skaalure, S.C., and Bryant, S.J. (2011). Gel structure has an impact on pericellular and extracellular matrix deposition, which subsequently alters metabolic activities in chondrocyte-laden PEG hydrogels. Acta Biomater. 7, 492–504. 10.1016/j.actbio.2010.08.021.

22. Bryant, S.J., and Anseth, K.S. (2002). Hydrogel properties influence ECM production by chondrocytes photoencapsulated in poly(ethylene glycol) hydrogels. J. Biomed. Mater. Res. 59, 63–72. 10.1002/jbm.1217.

23. Yan, X., Chen, Y.R., Song, Y.F., Ye, J., Yang, M., Xu, B.B., Zhang, J.Y., Wang, X., and Yu, J.K. (2020). Advances in the Application of Supramolecular Hydrogels for Stem Cell Delivery and Cartilage Tissue Engineering. Front. Bioeng. Biotechnol. 8, 1–13. 10.3389/fbioe.2020.00847.

24. Xu, J., Feng, Q., Lin, S., Yuan, W., Li, R., Li, J., Wei, K., Chen, X., Zhang, K., Yang, Y., et al. (2019). Injectable stem cell-laden supramolecular hydrogels enhance in situ osteochondral regeneration via the sustained co-delivery of hydrophilic and hydrophobic chondrogenic molecules. Biomaterials 210, 51–61. 10.1016/j.biomaterials.2019.04.031.

25. Ma, X., Yang, R., Wang, P., Liu, S., Xu, H., Ye, Z., and Chi, B. (2022). Bioinspired dual dynamic network hydrogels promote cartilage regeneration through regulating BMSC chondrogenic differentiation. Mater. Today Chem. 23. 10.1016/j.mtchem.2021.100648.

26. Stampoultzis, T., Guo, Y., Nasrollahzadeh, N., Karami, P., and Pioletti, D.P. (2022). Mimicking Loading-Induced Cartilage Self-Heating in Vitro Promotes Matrix Formation in Chondrocyte-Laden Constructs with Different Mechanical Properties. ACS Biomater. Sci. Eng. 10.1021/acsbiomaterials.2c00723.

27. Wei, K., Chen, X., Li, R., Feng, Q., and Bian, L. (2017). Multivalent Host-Guest Hydrogels as Fatigue-Resistant 3D Matrix for Excessive Mechanical Stimulation of Encapsulated Cells. Chem. Mater. 29, 8604–8610. 10.1021/acs.chemmater.7b02196.

28. Tong, Z., Jin, L., Oliveira, J.M., Reis, R.L., Zhong, Q., Mao, Z., and Gao, C. (2021). Adaptable hydrogel with reversible linkages for regenerative medicine: Dynamic mechanical microenvironment for cells. Bioact. Mater. 6, 1375–1387. 10.1016/j.bioactmat.2020.10.029.

