## Supporting Information for "Impact of Molecular Dynamics of Polyrotaxanes on Chondrocytes in Double Network Supramolecular Hydrogels under Physiological Thermomechanical Stimulation"

Author information:

1. Theofanis Stampoultzis

2. Vijay Kumar Rana

3. Yanheng Guo

4. Dominique P. Pioletti

**Effect of pure PEG and pure α-CD on gene expression**

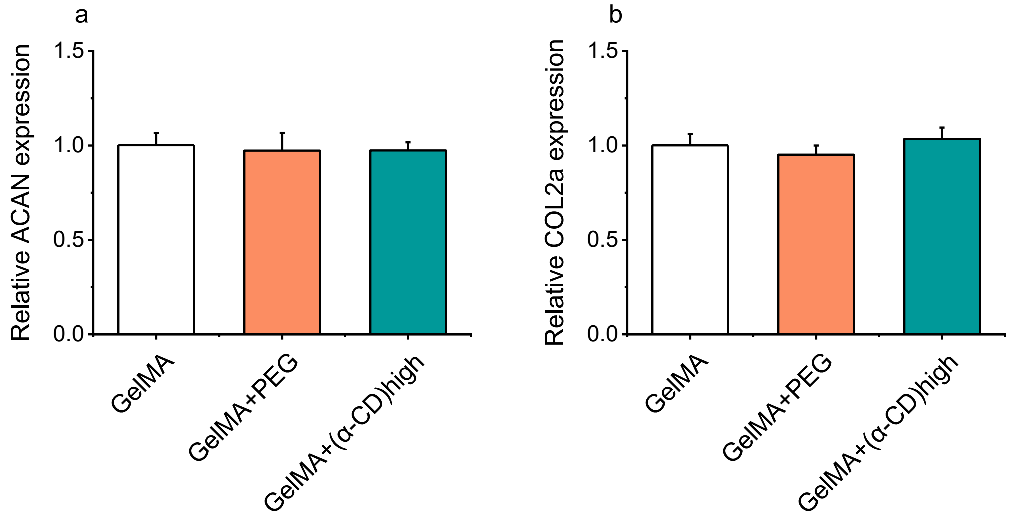


Figure S1 – Relative expression of genes of interest to assess the effects of individual PEG chains and α-CD molecules.

**Energy dissipation level of different hydrogel formulations**

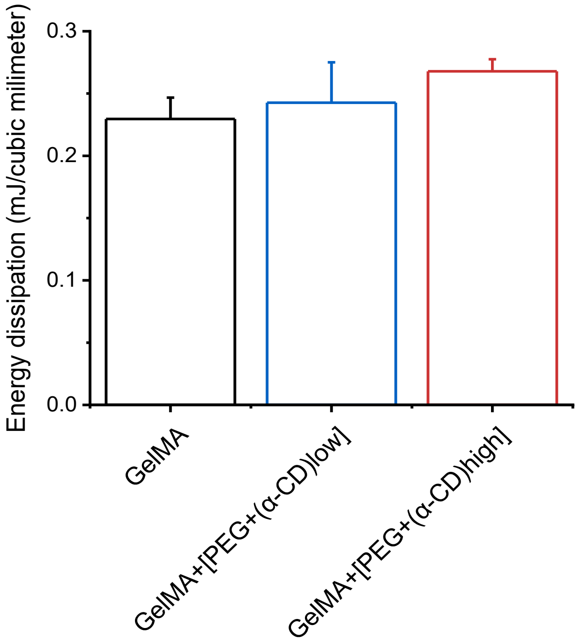


Figure S2 – Energy dissipation of hydrogel formulations calculated from the load-displacement curves of each sample. No significant differences were detected.
